# Plants with promising antileishmanial activity in Colombia: a systematic review and meta-analysis

**DOI:** 10.1101/2025.07.20.665797

**Authors:** C. Nieto-Clavijo, L. Morales, G. Zambrano, A. Delgado-Aldana, Z. Corredor-Rozo, EP. Calvo, D. Tinjacá, J. Chaparro-Olaya

## Abstract

**Introduction:** Leishmaniasis remains a major public health challenge in Colombia, driven by high incidence, *Leishmania* species diversity, and drug resistance. Colombian medicinal plants, rooted in rich ethnobotanical traditions and supported by the country’s exceptional biodiversity, represent a promising yet underexplored resource for the development of novel antileishmanial therapies.

**Aim of the study:** To systematically assess Colombian medicinal plants with reported *in vitro* antileishmanial activity, to estimate pooled IC_50_ values through meta-analysis, and to identify extracts with favorable selectivity indices (SI) as potential antileishmanial candidates.

**Materials and methods:** A systematic search (2000–April 2025) of PubMed, EMBASE, and LILACS identified *in vitro* studies reporting IC_50_ values of Colombian plant extracts against *Leishmania* spp. A random-effects meta-analysis was used to estimate pooled IC_50_ values. Risk of bias was assessed using a modified QUIN tool. Subgroup analyses explored methodological and biological factors, such as plant part, extraction solvent, and taxonomic family.

**Results:** Twenty-five studies were included, covering 110 records on 24 plant species. Thirteen studies with complete IC_50_ ± SD data were eligible for meta-analysis. The pooled mean IC_50_ was 41.25 µg/mL (95% CI: 37.95–44.55), with substantial heterogeneity (I^2^ = 100%). Lower IC_50_ values were associated with bark/wood extracts, methanol or dichloromethane solvents, and species from the Chrysobalanaceae and Bignoniaceae families. *Xylopia discreta* (Annonaceae) showed the highest SI (up to 110). Additional species from the Lamiaceae, Lauraceae, Picramniaceae, and Rutaceae families also demonstrated favorable SI values. However, methodological variability limited the ability to make direct comparisons across studies.

**Conclusion:** Several Colombian plant species showed promising *in vitro* antileishmanial activity, with selected extracts combining high potency and notably high selectivity. The inclusion of SI analysis in this review provides a more meaningful assessment of therapeutic potential beyond IC_50_ values alone. These findings underscore the value of Colombia’s plant biodiversity as a source of candidate compounds for antileishmanial drug development. However, standardized *in vitro* protocols, consistent cytotoxicity evaluation, and *in vivo* validation remain essential to ensure comparability and guide the selection and advancement of the most promising extracts toward therapeutic application.

## Introduction

Leishmaniasis is a neglected tropical disease caused by protozoan parasites of the genus *Leishmania*, transmitted to humans through the bite of infected female *Phlebotomine* sandflies (*Phlebotomus* spp. in the Old World and *Lutzomyia* spp. in the New World) [1]. This vector-borne disease presents a wide clinical spectrum, with three main manifestations: cutaneous leishmaniasis (CL), the most prevalent form, characterized by ulcerative skin lesions that may result in disfiguring scars; mucocutaneous leishmaniasis (MCL), a severe and potentially mutilating progression of CL that affects the mucous membranes of the nasopharyngeal region; and visceral leishmaniasis (VL), the most severe form, which involves internal organs such as the spleen, liver, and bone marrow, leading to systemic complications and high mortality if left untreated [2, 3].

In Colombia, nine *Leishmania* species have been historically reported: *L. panamensis*, *L. braziliensis*, *L. guyanensis*, *L. equatoriensis*, *L. lainsoni*, *L. colombiensis*, *L. mexicana*, *L. amazonensis*, and *L. infantum*. All except *L. infantum*, which is associated with VL, have been linked to CL. Furthermore, *L. panamensis*, *L. braziliensis*, and *L. guyanensis* have also been associated with MCL [4]. In addition, *L. naiffi* and *L. lindenbergi* were identified more recently in three Colombian soldiers, representing the first reports of these species in the country [5]. Compared to 2023, an overall increase of 25.5% in reported cases of leishmaniasis in Colombia was observed in 2024, from 5,486 to 6,886 notifications [6, 7]. This increase was mainly driven by a 26.2% increase in CL, which continues to represent the most frequent clinical manifestation. In addition, a moderate increase of 6.6% was observed in cases of MCL, while cases of VL decreased by 15% [7]. The persistent epidemiological burden of leishmaniasis in Colombia, driven by its endemicity, high incidence rates, broad geographical distribution, and the diversity of *Leishmania* species, highlights the urgent need for improved disease management Conventional treatment strategies for leishmaniasis in Colombia rely primarily on the use of pentavalent antimonials, amphotericin B, miltefosine, and other pharmacological agents [8]. Despite their widespread use, these therapies are associated with significant limitations, including severe toxicity, high cost, complex administration protocols, and, most critically, the increasing emergence of drug-resistant *Leishmania* strains. A recent systematic review addressing treatment failure and clinical relapse in leishmaniasis identified Latin America as one of the regions with the highest number of cases, with Brazil, Colombia, and French Guiana reporting the highest rates of relapse [9]. The study also highlighted the impact of treatment failure across clinical forms of the disease, reporting failure or relapse rates of 47.6% for CL and 45.2% for VL. These findings underscore the urgent need for novel therapeutic strategies capable of overcoming the challenges posed by drug-resistant *Leishmania* strains.

Within the repertoire of pharmacological approaches, increasing attention is being paid to the reuse of existing drugs and the identification of new compounds with anti-leishmanial activity. In this context, natural products have emerged as a particularly promising area of exploration. Medicinal plants offer a rich source of bioactive molecules that may lead to the development of safer, more effective, and affordable therapeutic options, addressing key gaps in current pharmacological interventions.

Colombia’s exceptional biodiversity presents a valuable opportunity for bioprospecting efforts. As one of the most biodiverse countries in the world, Colombia hosts a wide range of plant species, many of which produce secondary metabolites with antiparasitic activity [10]. This phytochemical diversity constitutes a largely untapped resource for the discovery of new leishmanicidal compounds. Recent pharmacological and ethnobotanical studies have identified several species with activity against *Leishmania* spp. [11], further supporting the need for a systematic evaluation of the available evidence.

This systematic review and meta-analysis aims to comprehensively analyze research conducted between January 1, 2000, and April 30, 2025, on Colombian medicinal plants with documented antileishmanial activity. By synthesizing the available evidence, the study aims to identify candidate species and their bioactive constituents with potential for the development of safer and more effective treatments for leishmaniasis. Additionally, it aims to identify current knowledge gaps and propose future research priorities to support the advancement of these natural products toward clinical application.

## Methodology

### Reporting System and Registration

This systematic review and meta-analysis was conducted in accordance with the Preferred Reporting Items for Systematic Reviews and Meta-Analyses (PRISMA) [12]. The study protocol was prospectively registered in the International Prospective Register of Systematic Reviews (PROSPERO) under the registration number CRD420251076470, accessible at https://www.crd.york.ac.uk/PROSPERO/view/CRD420251076470.

### Data Sources

A comprehensive literature search was conducted in the PubMed, EMBASE, and LILACS databases. These platforms were selected for their complementary coverage of biomedical, pharmacological, and regional scientific literature. The search was restricted to articles published between January 1, 2000, and April 30, 2025, in English, Spanish, or Portuguese.

### Search Strategy

Search queries combined Medical Subject Headings (MeSH) and free-text terms, linked with Boolean operators (“AND” and “OR”) to optimize sensitivity and specificity. The full search strategies for each database are provided in the Supporting information (S1 Table). An information specialist contributed to the development and refinement of search terms. We followed the PRISMA 2020 checklist for reporting (S2 Table).

### Eligibility criteria

Studies were included if they (i) reported *in vitro* leishmanicidal activity of Colombian plant materials, (ii) provided IC_50_ values for antiparasitic activity, (iii) were published as full-text articles, and (iv) were written in English, Spanish, or Portuguese. Studies were excluded if they (i) lacked IC_50_ data, (ii) did not clearly confirm the Colombian origin of the plant material or failed to cite a primary source verifying it, (iii) were not available in full-text format, (iv) consisted of publication types such as conference abstracts, reviews, letters to the editor, posters, infographics, or any other format lacking complete data, (v) relied exclusively on *in silico* approaches, or (vi) evaluated only synthetic derivatives.

### Study Selection and Quality Assessment

Study selection was conducted by three independent reviewers who screened titles, abstracts, and full texts according to the predefined eligibility criteria. Discrepancies were resolved by consensus. For quality assessment, a modified version of the quality assessment tool for *in vitro* Studies (QUIN tool) [13] was applied. Each study was evaluated based on specific methodological components, scoring 2 points for clearly and adequately reported elements, 1 point for insufficiently specified ones, and 0 points for those not reported. The total score was expressed as a percentage of the maximum possible, and studies were classified into three quality levels: poor (<50%), moderate (50–75%), or high (>75%).

### Statistical analysis

To estimate the pooled inhibitory effect of plant extracts and isolated compounds against *Leishmania* spp., a meta-analysis was conducted by calculating the mean IC_50_ values and their corresponding 95% confidence intervals (CIs) from the included studies. Given the anticipated heterogeneity across studies, a random-effects model was applied. Forest plots were generated to visualize both individual and pooled IC_50_ estimates along with their CIs.

Between-study heterogeneity was assessed using Cochran’s Q test (with *p* < 0.1 indicating significant heterogeneity) and the I^2^ statistic, interpreted as low (25%–49%), moderate (50%– 74%), or high (≥75%). Potential publication bias was evaluated using funnel plots and Egger’s test. All statistical analyses were performed using Stata 16.0 software (StataCorp, College Station, TX, USA).

## Results

### Study selection and data classification

A total of 115 articles were initially retrieved through database searches. After eliminating 27 duplicates, the titles and abstracts of 88 studies were examined. Of these, 55 were excluded because they were reviews, did not refer to plant species collected in Colombia, or did not focus on antileishmanial biological activity. The remaining 33 articles were evaluated in full text. Eight additional studies were excluded due to lack of access to the full text, inconsistent data, use of non-Colombian plant material, exclusive application of *in silico* models, or for focusing only on synthetic derivatives. Finally, 25 studies met the inclusion criteria and were retained for analysis (Figure 1).

**Figure 1.**
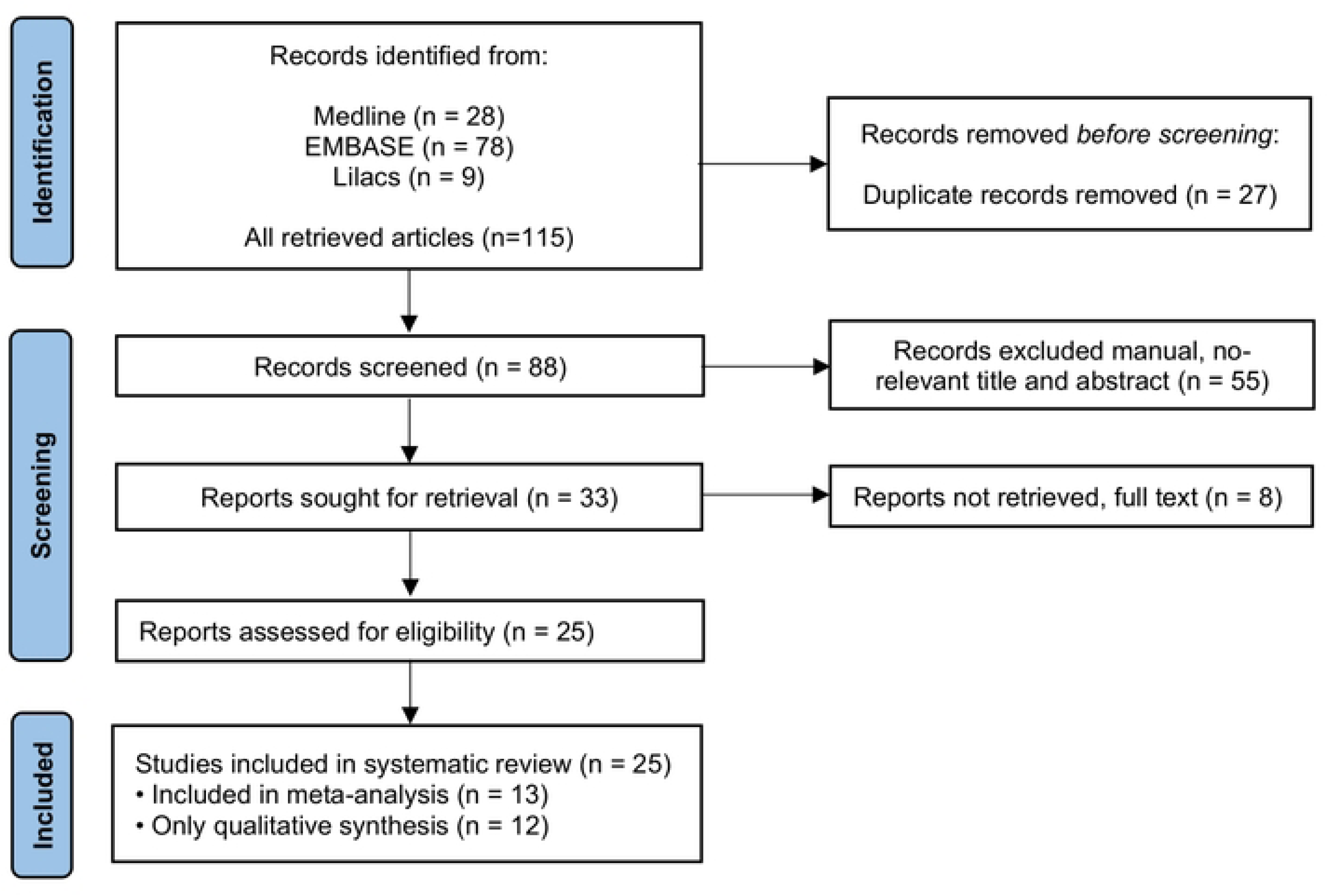
Flowchart describing the study design process.

Although this review aimed to evaluate antileishmanial efficacy based on IC_50_ values, the included studies varied considerably in terms of data quality and completeness. Thirteen studies met the criteria for inclusion in the meta-analysis by reporting IC_50_ values along with measures of variability, such as standard deviations and the number of replicates. A second group of studies reported IC_50_ values but were excluded from the meta-analysis because the data were either outside the quantifiable range or lacked essential statistical parameters. Finally, a third group of studies did not provide quantitative IC_50_ values but presented qualitative assessments of leishmanicidal activity, classifying it as “low,” “moderate,” or “high,” or describing it narratively. While these limitations precluded meta-analysis and the construction of forest plots for the latter two groups, the data still contribute valuable insights for identifying promising species, informing future research priorities, and avoiding duplication of efforts. To enhance the comparability and utility of future studies, we recommend the standardized reporting of efficacy metrics alongside appropriate measures of variability.

### Risk of bias assessment

To assess methodological quality and reporting standards, the included studies were evaluated using a modified QUIN tool for *in vitro* research (S3 Table). The results are summarized in Figure 2. Most of the included studies demonstrated adequate methodological clarity across most assessed domains, particularly in the presentation of results, outcome measurement methods, and the overall description of the experimental procedures. However, the domain with the most frequent shortcomings was statistical analysis. A substantial proportion of studies (about 20%) lacked sufficient detail, and nearly 30% did not specify the statistical procedures at all. This indicates that, although quantitative data were available, many studies provided limited detail on how variability, statistical significance, or effect sizes were assessed, potentially compromising the reproducibility and comparability of their findings. Additional methodological limitations were noted in the reporting of sampling techniques and the clarity of study aims or objectives. These elements were inadequately described or omitted in 25% and 20% of the studies, respectively. Incomplete reporting of sampling strategies is particularly relevant in plant-based studies, where ecological variability and harvesting protocols can influence phytochemical content and, consequently, bioactivity outcomes. In contrast, over 90% of the studies clearly described the comparison groups and methodological procedures, suggesting a generally robust experimental design and implementation, despite shortcomings in other reporting areas.

**Figure 2.**
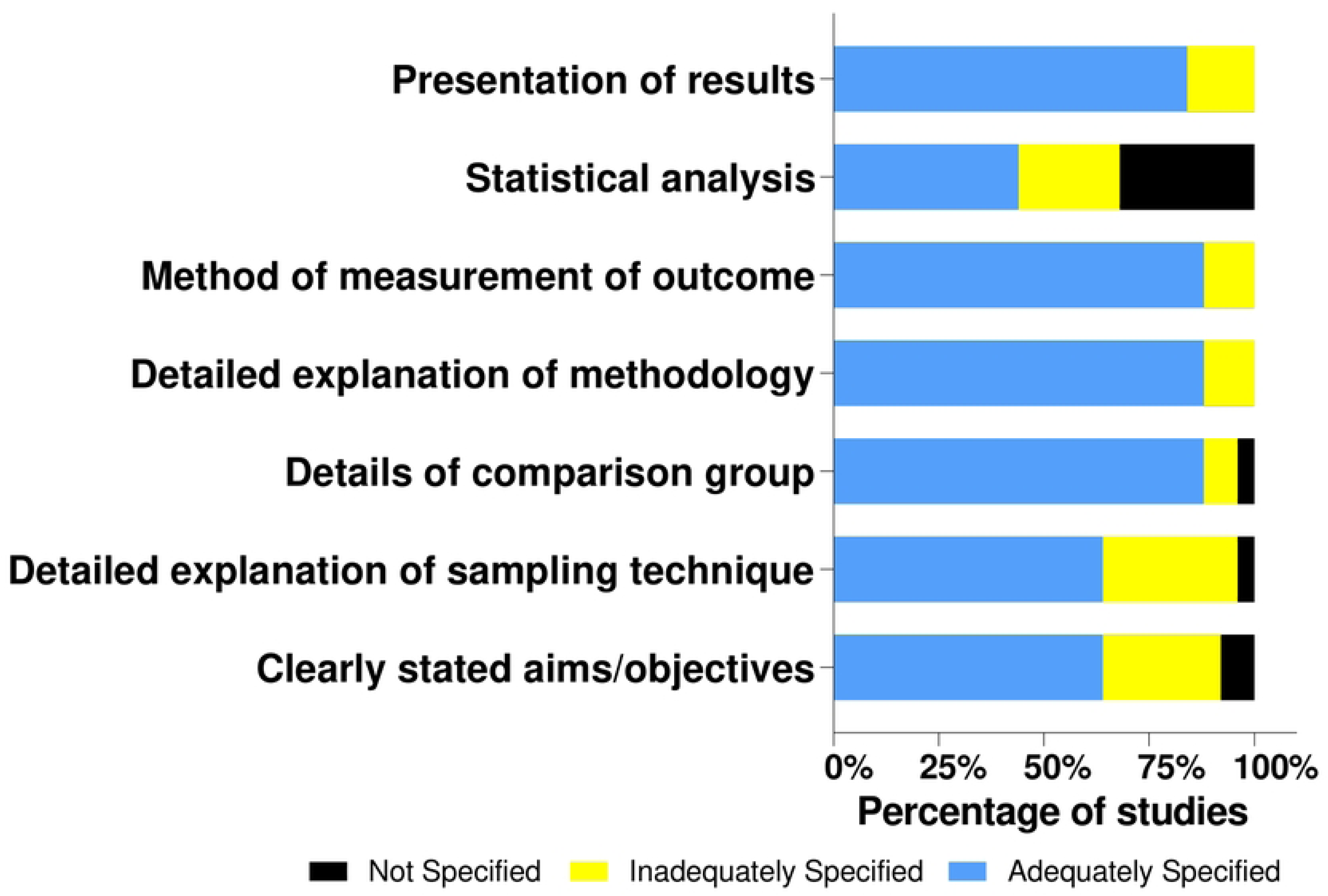
Methodological quality across six key domains in the included studies. The y-axis lists the evaluated methodological domains. The x-axis represents the percentage of studies assessed. Bars are stacked and divided into three colors indicating the level of reporting: Blue = adequately reported, yellow = partially reported, and black = not reported. Each bar sums to 100% and shows the proportion of studies falling into each reporting category per domain.

These findings highlight the need for greater adherence to standardized reporting guidelines in *in vitro* pharmacological research, to improve clarity, reproducibility, and quality assessment of preclinical evidence derived from natural products.

### Characterization of bioactive extracts and experimental models

From the 13 studies included in the meta-analysis, 110 records were extracted corresponding to different extracts, fractions or metabolites from 24 plant species collected in Colombia and evaluated *in vitro* against various species of *Leishmania*. The most frequently studied species was *L. panamensis* (64/110), followed by *L. braziliensis* (25/110), *L. major* (9/110), *L. donovani* (4/110), *L. amazonensis* (4/110) and *L. guyanensis* (4/110). The promastigote stage was the most commonly used (68/110) (Table 1 & 2).

**Table 1.**
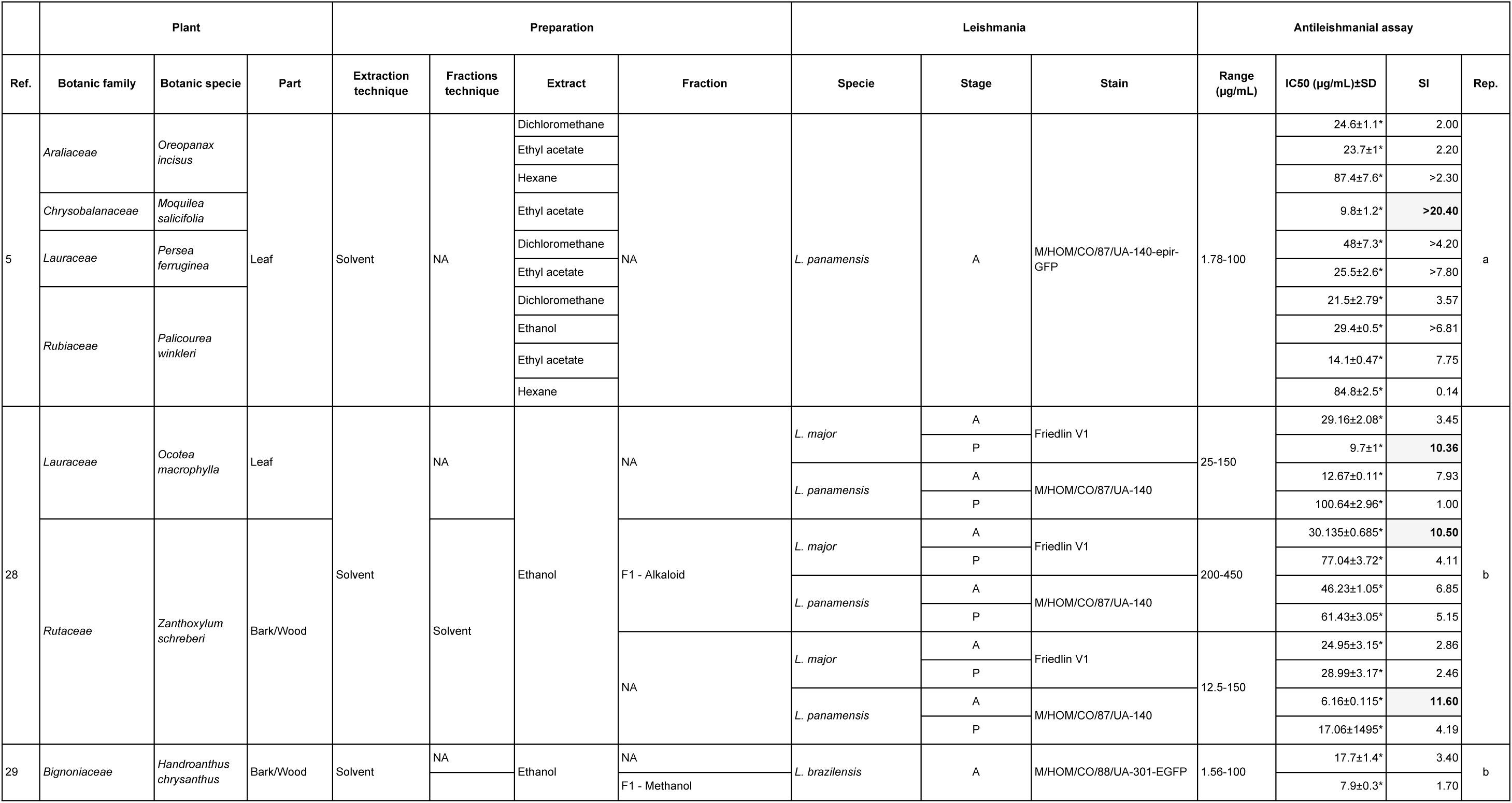
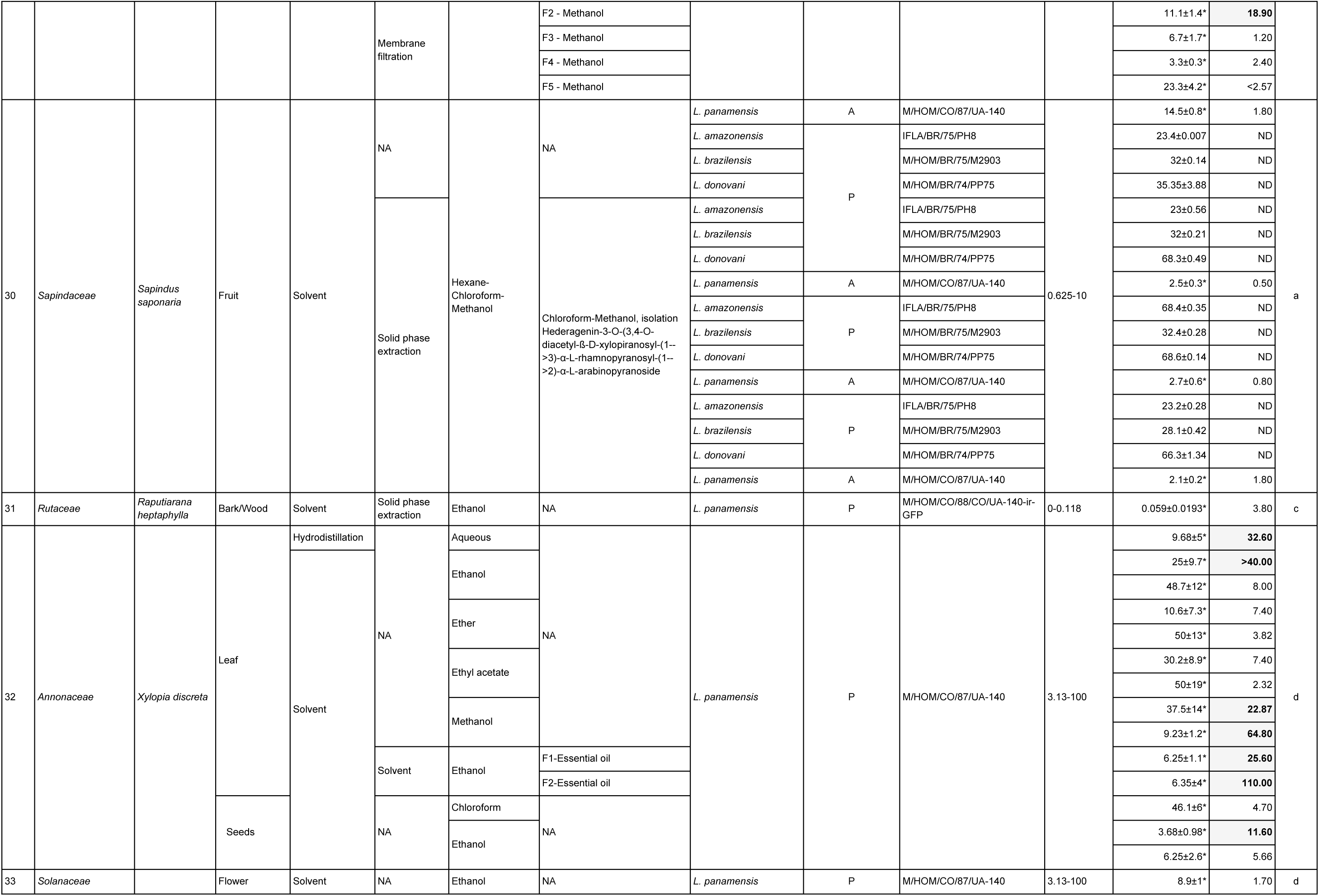
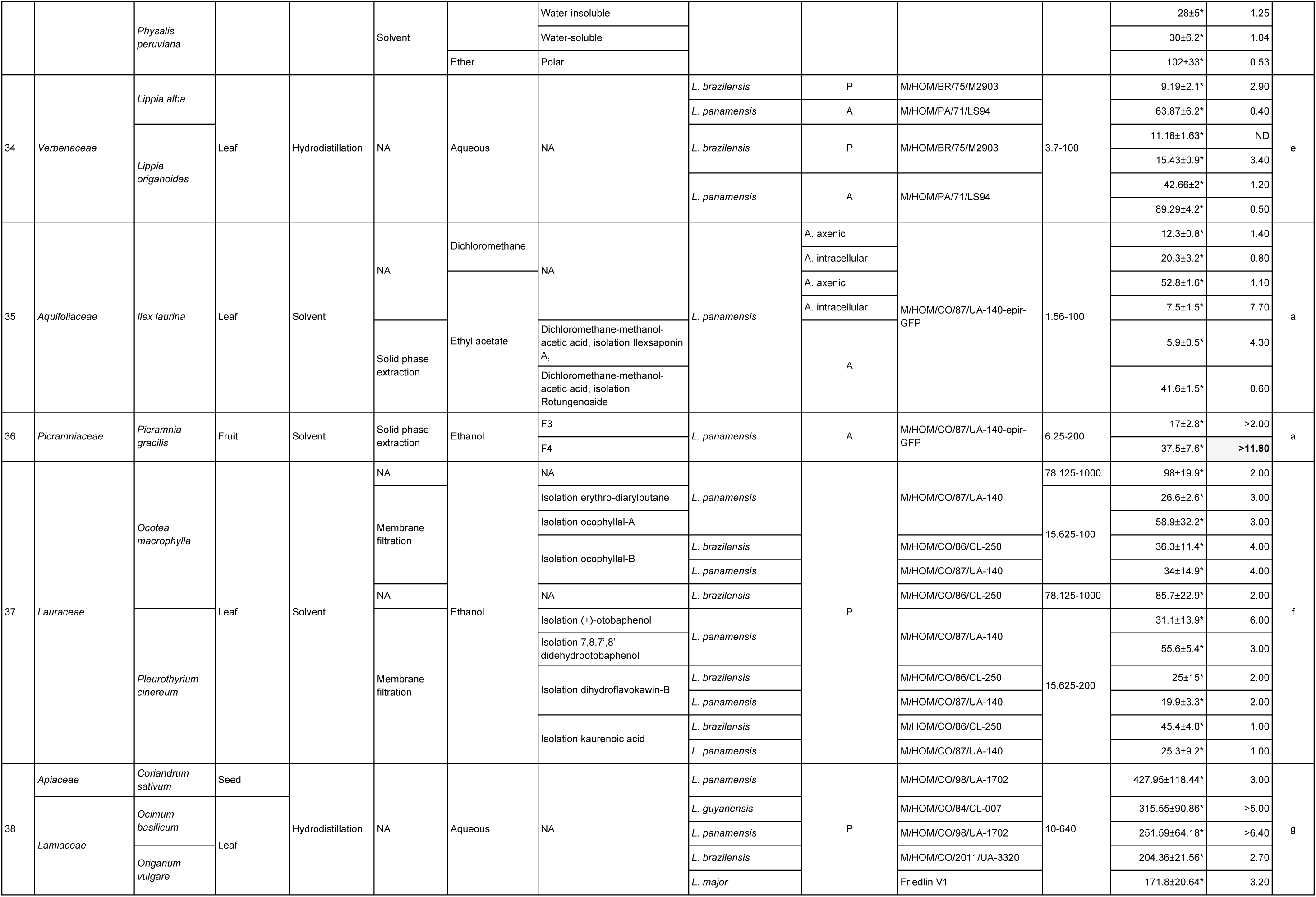
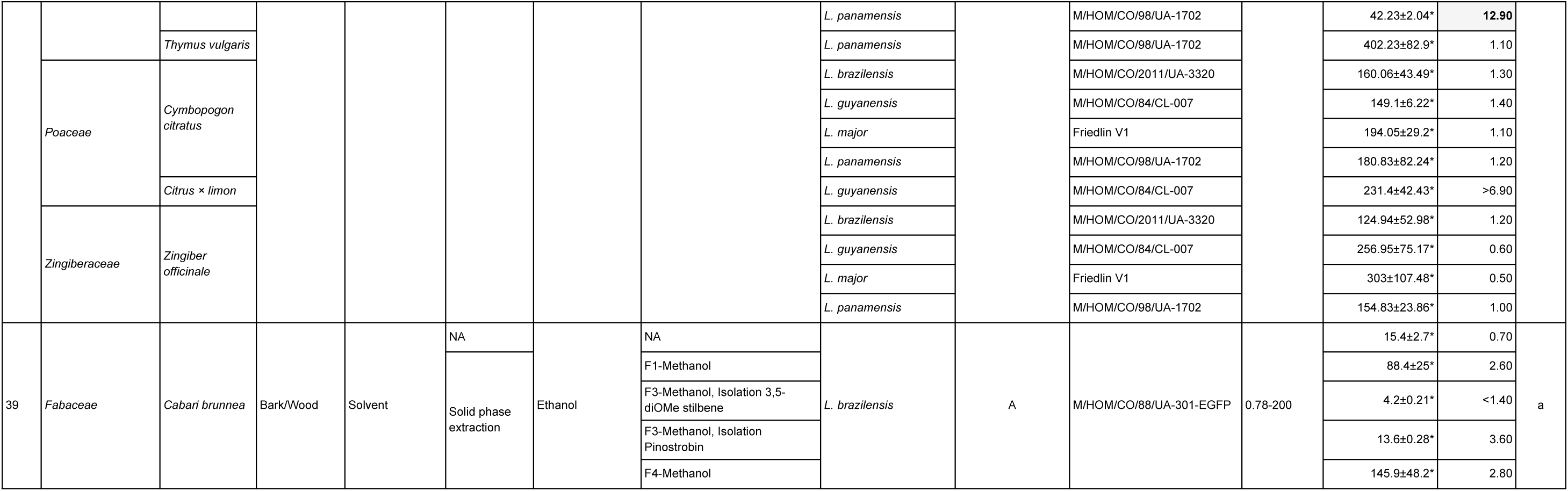
Summary of plant species, parasite strains, and reported antileishmanial activity in studies included in the meta-analysis. IC_50_ refers to the concentration required to inhibit 50% of parasite viability. In cases marked with an asterisk (*), inhibition was measured using EC_50_. The selectivity index (SI), calculated as the ratio of cytotoxic concentration to effective antiparasitic concentration, provides an estimate of therapeutic potential. SI values greater than 10 are shown in bold within shaded cells, as this threshold is commonly considered a minimum criterion for preclinical evaluation, indicating reduced toxicity to mammalian cells. NA, not applicable; ND, not determined; A, amastigote; P, promastigote. Superscript letters denote experimental conditions as follows: a: two independent trials with triplicates; b: triplicate; c: five independent experiments; d: three independent trials with triplicates; e: duplicate; f: three independent trials with duplicates; g: two independent trials with duplicates.

Most extracts were obtained from species belonging to the families Lauraceae (n = 18), Sapindaceae (n = 15), and Annonaceae (n = 14). *Sapindus saponaria* was the most frequently studied species (n = 16), with reported traditional uses for treating leishmaniasis-related ulcers, as well as gastrointestinal and inflammatory disorders. *Xylopia discreta* was also frequently cited (n = 14), although no traditional uses were mentioned in the reviewed studies. Leaves were the most commonly used plant part (n = 64), followed by bark/wood (n = 20), fruits (n = 18), seeds (n = 4), and flowers (n = 4). Regarding extraction solvents, ethanol was the most frequently used (n = 48). The characteristics of the bioactive extracts are summarized in Tables 1 and 2.

**Table 2.**
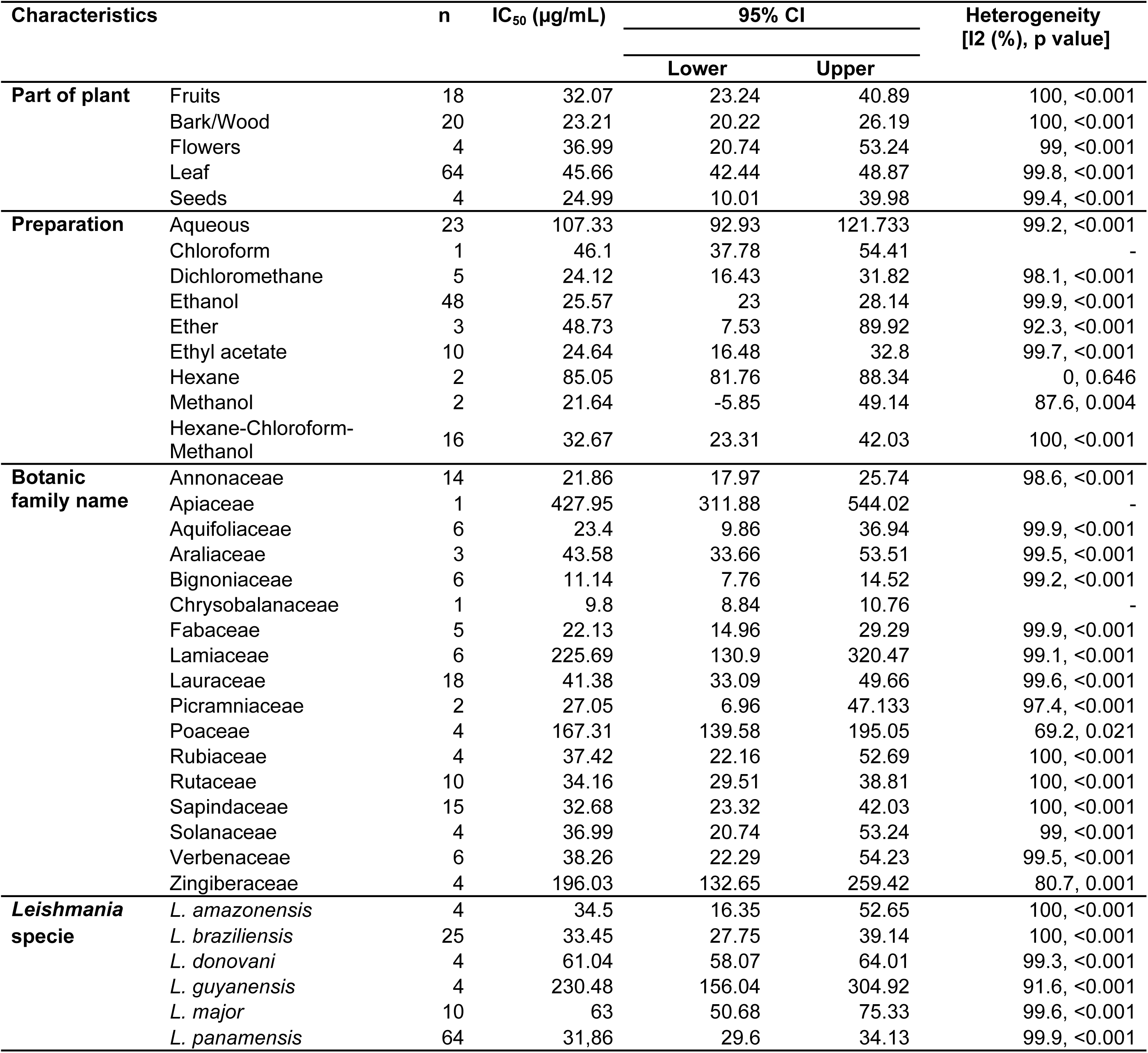
Subgroup analyses of antileishmanial activity by plant part, extraction method, botanical family, and leishmania species. Summary of subgroup analyses from 13 studies included in the meta-analysis of plant-derived extracts with reported IC_50_ values. Data are presented as pooled mean IC_50_ values (µg/mL) with 95% confidence intervals (CI). Subgroups are stratified by plant part, preparation method, botanical family, and Leishmania species. Heterogeneity is reported as I^2^ (%) and associated p value. n indicates the number of records contributing to each estimate. Heterogeneity statistics are not provided for subgroups represented by a single study.

### IC50 and selectivity index values

It is important to note that a low IC_50_ indicates biological activity but does not necessarily imply selectivity. The selectivity index (SI), defined as the ratio between the cytotoxic concentration to host cells (CC_50_) and the inhibitory concentration against *Leishmania* spp (IC_50_), is a key parameter for evaluating how selectively a compound targets the parasite without harming host cells. Compounds with moderate or relatively high IC_50_ values but elevated SI values may represent safer and more selective candidates, whereas those with low IC_50_ values but no cytotoxicity data cannot be adequately assessed. Unfortunately, a meta-analysis or comparative evaluation based on SI was not feasible in this review, as 26 of the 110 records did not report SI values. Moreover, among the studies that did report SI, only a few provided associated measures of variability.

According to the literature, extracts with SI ≥ 10 are considered selective and suitable for further studies, while values above 20 indicate high selectivity and therapeutic potential [14, 15]. Among the selective extracts identified, those tested against *L. panamensis* promastigotes exhibited the highest SI. The most selective extract was obtained from *X. discreta* (SI = 110.0), followed by *Origanum vulgare* (SI = 12.90), both exceeding the established threshold for potential further investigation. The same for the amastigote form of *L. panamensis*, for which, the most favorable selectivity was observed for extracts from *Moquilea salicifolia* (SI >20.40), *Picramnia gracilis* (SI >11.80), and *Zanthoxylum schreberi* (SI = 11.60). In the case of *L. braziliensis* amastigotes, *Handroanthus chrysanthus* demonstrated high selectivity (SI = 18.90), approaching the threshold indicative of strong therapeutic potential. For *L. major*, selectivity was observed for *Z. schreberi* (SI = 10.50, amastigote stage) and *Ocotea macrophylla* (SI = 10.36, promastigote stage), suggesting that these extracts may also merit further evaluation. Notably, *X. discreta* consistently demonstrated high selectivity indices across different extraction methods. Whether obtained through methanolic, ethanolic, aqueous, or essential oil preparations, the extracts exhibited SI values well above the threshold for further investigation, highlighting the robustness of its leishmanicidal potential. This consistency across extraction types positions *X. discreta* as a particularly promising candidate for anti-*Leishmania* drug development (Table 1).

### Meta-analysis

Regarding IC_50_ data, heterogeneity analysis revealed substantial variability among studies (Q = 7.8 × 10⁶, df = 109, I^2^ = 100%, p < 0.05), warranting the use of a random-effects model (Figure 3). The pooled mean IC_50_ was 41.25 µg/mL (95% CI: 37.95 – 44.55). Publication bias was assessed using a funnel plot and Egger’s regression test. Visual inspection of the funnel plot (Figure 4) suggested asymmetry, with studies concentrated in the upper right quadrant and sparse in the lower left, indicative of potential reporting bias. Nevertheless, Egger’s test did not detect significant publication bias (t = –0.25, p = 0.804). Subgroup analyses revealed considerable heterogeneity in leishmanicidal activity according to the plant part used, type of extract, botanical family, and *Leishmania* species. Most subgroups showed I^2^ values greater than 90% and p-values < 0.001 (Cochran’s Q test), indicating significant differences among studies. This high degree of variability can be attributed to the considerable differences in the overall experimental systems across studies. These include variation in the plant species used, the geographical location of plant collection, the specific plant part tested, and the preparation method of the extracts, ranging from crude extracts to fractions and purified metabolites. Additionally, heterogeneity arose from differences in the *Leishmania* species and strains employed for *in vitro* testing, as well as in broader methodological and experimental conditions. These findings underscore the critical need to standardize protocols in future studies to facilitate more robust and meaningful comparisons. Regarding the plant part used, leaf extracts exhibited the highest mean IC_50_ (45.66 µg/mL; 95% CI: 42.44–48.87), whereas bark/wood extracts showed lower values (23.21 µg/mL; 95% CI: 20.22–26.19). All subgroups presented high heterogeneity (I^2^ > 99%; p < 0.001). Among solvents, aqueous extracts had the highest mean IC_50_ (107.33 µg/mL; 95% CI: 92.93–121.73), while ethanol (25.57 µg/mL; 95% CI: 23.00 – 28.14) and dichloromethane extracts (24.12 µg/mL; 95% CI: 16.43 – 31.82) showed greater potency. Hexane extracts were the only subgroup with low heterogeneity (I^2^ = 0%; p = 0.646). The botanical families with the lowest mean IC_50_ values were Bignoniaceae (11.14 µg/mL; 95% CI: 7.76 – 14.52) and Chrysobalanaceae (9.80 µg/mL; 95% CI: 8.84–10.76), whereas Lamiaceae (225.69 µg/mL; 95% CI: 130.90 – 320.47) and Zingiberaceae (196.03 µg/mL; 95% CI: 132.65 – 259.42) recorded the highest values. Finally, pooled mean IC_50_ values ranged from 31.86 µg/mL for *L. panamensis* (95% CI: 29.60 – 34.13) to 230.48 µg/mL for *L. guyanensis* (95% CI: 156.04 – 304.92), with high heterogeneity across all species subgroups (I^2^ > 99%; p < 0.001). Detailed strain-specific data are presented in Table 2.

**Figure 3.**
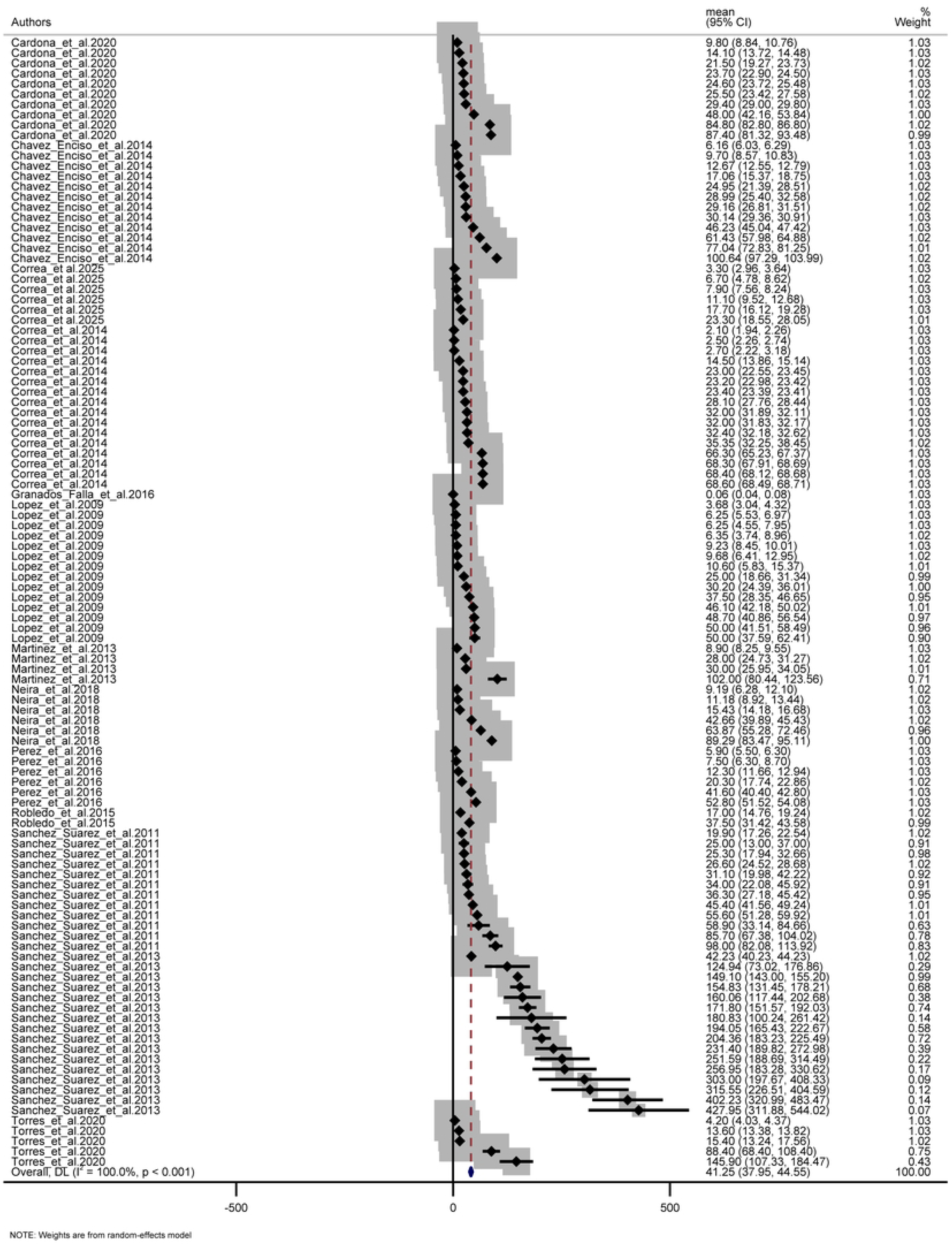
Forest Plot from Random-Effects Meta-Analysis of the mean IC_50_ values of medicinal plants with promising antileishmanial activity in Colombia. The pooled mean IC_50_ was 41.25 µg/mL (95% CI: 37.95–44.55). Heterogeneity across studies was assessed using Cochran’s Q test (Q = 7.8 × 10⁶, df = 109, p < 0.05) and the I^2^ statistic (I^2^ = 100.0%, p < 0.001), indicating substantial between-study variability.

**Figure 4.**
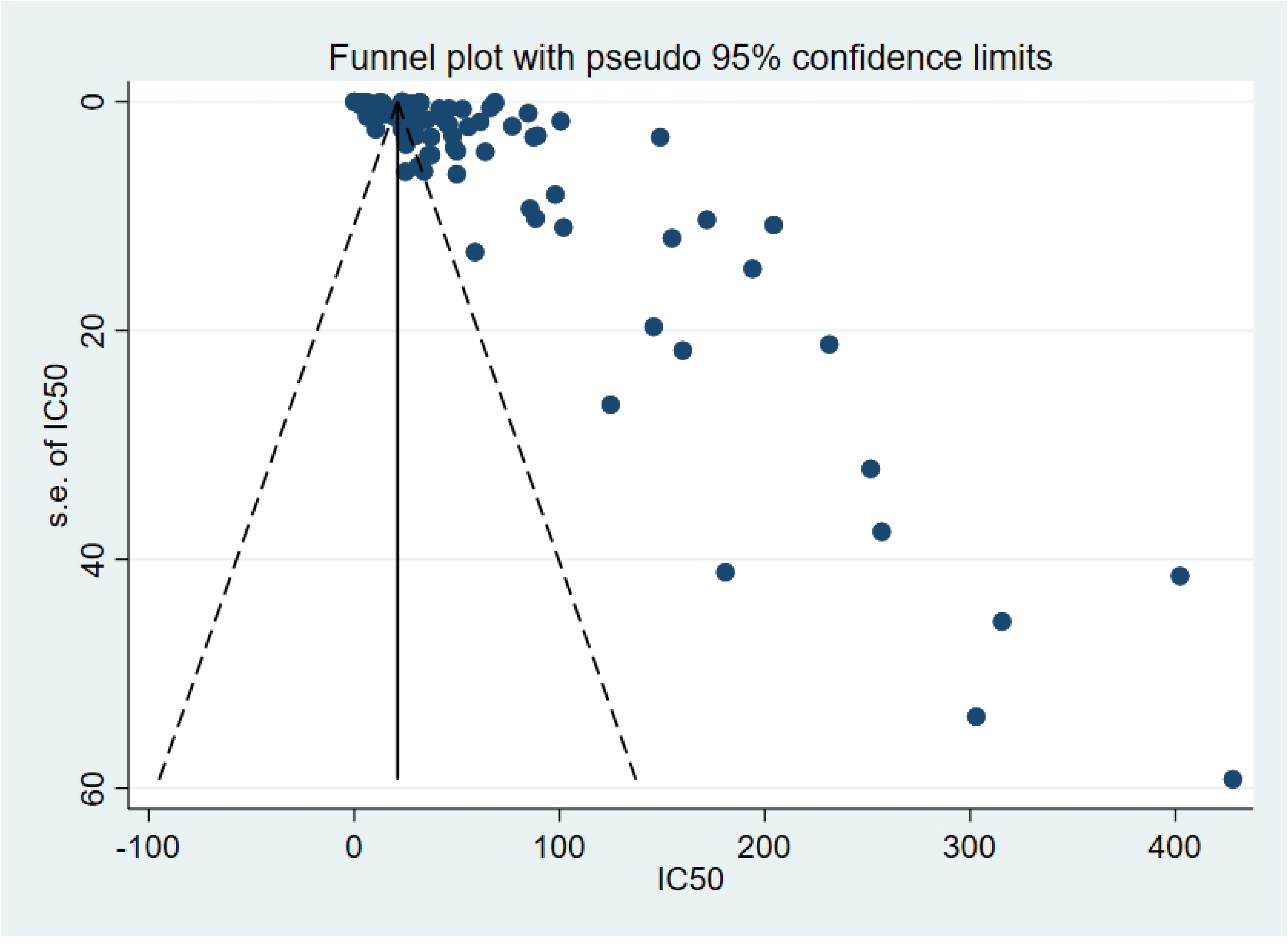
Funnel Plot of Studies Included in the Meta-Analysis of IC_50_ values of medicinal plants collected in Colombia. Funnel plot used to assess publication bias among the studies included in the meta-analysis of IC_50_ values. Each point represents an individual study, with the x-axis indicating the reported IC_50_ (µg/mL) and the y-axis representing the standard error of the estimate.

### Qualitative and semi-quantitative data

As previously noted, in addition to the quantitative analysis, records containing both qualitative and semi-quantitative data on leishmanicidal activity were included. These correspond to extracts for which exact IC_50_ values were not reported either because they fell outside the measurable range of the assay or were assessed solely through descriptive methods. This information is summarized in figure 5, which categorizes the extracts based on their IC_50_ values (≤10 µg/mL; 10–25 µg/mL; >25–50 µg/mL; >50–100 µg/mL; and >100 µg/mL). This visual representation facilitates the rapid identification of extracts with potentially relevant activity, even in the absence of complete quantitative data. Among these, promising extracts were observed from genera such as *Xylopia*, *Annona*, *Sapindus*, *Lippia*, and *Croton*, which may guide future research toward more rigorous evaluation, including toxicity and selectivity assessments as well as phytochemical characterization.

**Figure 5.**
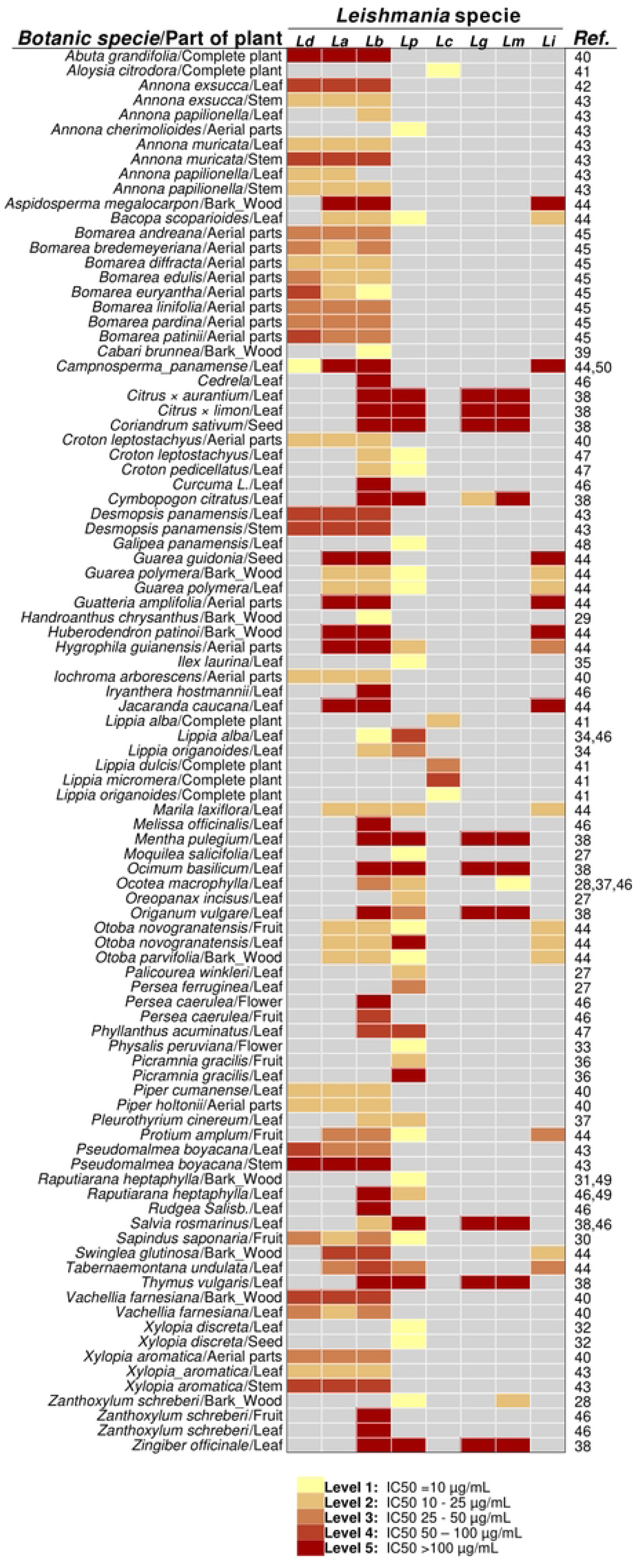
Heat Map of Leishmanicidal Activity (IC_50_) of Plant Extracts by Plant Part and *Leishmania* Species. Each row represents a specific plant species and the part of the plant used for extract preparation, while columns correspond to different Leishmania species. Cell values indicate IC_50_ values (µg/mL) obtained from in vitro assays. Gray-shaded cells indicate combinations for which no data were reported in the studies included in the meta-analysis. Abbreviations: Ld, *L. donovani*; La, *L. amazonensis*; Lb, *L. braziliensis*; Lp, *L. panamensis*; Lc, *L. chagasi*; Lg, *L. guyanensis*; Lm, *L. major*; Li, *L. infantum*.

In the study by Calderón et al. (2010), ethanolic extracts from numerous plant species (collected across seven Latin American countries, including Colombia) were tested *in vitro* against *L. mexicana*. Although the authors reported that several extracts showed antileishmanial activity with IC_50_ values ≤ 50 µg/mL, none of the extracts derived from Colombian plant species fell within this threshold (S4 Table). All tested Colombian extracts exhibited IC_50_ values > 50 µg/mL and were therefore considered inactive or of no further interest by the authors. These findings still represent relevant preliminary background, as they demonstrate that extracts from those Colombian species were challenged in antileishmanial assays and yielded only modest activity.

## Discussion

The results of this systematic review and meta-analysis demonstrate that several plant species collected in Colombia exhibit promising *in vitro* antileishmanial activity, as reflected by IC_50_ values comparable to, or in some cases lower than, those reported in previous systematic reviews of plants collected in other countries. The pooled mean IC_50_ for antileishmanial plant extracts from Colombia was 41.25 µg/mL (95% CI: 37.95–44.55), notably lower than the pooled mean reported for herbal extracts in Iran, which was 456.64 µg/mL (95% CI: 396.15–517.12) [16]. In comparison, the pooled mean IC_50_ for Ethiopian medicinal plants was 16.80 µg/mL (95% CI: 12.44–21.16) against promastigotes and 13.81 µg/mL (95% CI: 13.12–14.50) against amastigotes [17]. While the IC_50_ is a key parameter for estimating the biological potency of an extract or compound, it is not sufficient to determine its viability as a pharmacological candidate. *In vitro* efficacy must be evaluated alongside toxicity and selectivity profiles. One of the main contributions of this review is the inclusion and analysis of the selectivity index (SI), which provides critical insight into the therapeutic window of a given extract. An extract with moderate IC_50_ but high SI may, in fact, be more promising than one with a lower IC_50_ but poor selectivity. Cytotoxicity and selectivity data are therefore essential for assessing biological specificity and guiding preclinical development of antileishmanial agents. These parameters should be considered fundamental benchmarks in future efforts to identify and validate novel therapeutic candidates.

In some notable cases, such as *X. discreta*, SI values exceeding 20 (up to 110) were reported, suggesting a highly favorable therapeutic window, particularly in the essential oil fraction tested against promastigotes of *L. panamensis*. This pattern of leishmanicidal activity combined with low cytotoxicity does not appear to be exclusive to *X. discreta* within the genus, as bioactive compounds with similar effects have been reported in other *Xylopia* species. For example, *X. parviflora* roots collected in Limpopo Province, South Africa, and extracted with dichloromethane inhibited the growth of *L. donovani* amastigotes with an IC_50_ of 5.01 μg/mL and an SI of 10 [18]. Likewise, a diterpene glycoside of the ent-kaurene type isolated from the leaves of *X. excellens* collected in Manaus, Brazil, exhibited *in vitro* activity against *L. amazonensis* promastigotes (IC_50_ = 15.23 ± 0.64 µg/mL) [19]. Although the authors considered the selectivity index of 1.96 to indicate good selectivity, this interpretation relies on a less conservative threshold and lacks supporting references. Nonetheless, these findings reinforce the relevance of the *Xylopia* genus as a promising source of antileishmanial metabolites and position the Colombian species *X. discreta*, with its consistently higher SI values, as a particularly compelling candidate for further investigation. These comparisons also underscore the importance of adopting standardized and evidence-based criteria for SI interpretation in order to facilitate the comparison of findings across studies and support the identification of truly promising therapeutic candidates.

The study by López et al. (2009) on *X. discreta* highlights the significant variability that can arise in both IC_50_ and selectivity index (SI) values depending on the type of extract used, offering a valuable opportunity to understand how extraction methods influence bioactivity profiles. In that study, a range of solvents (water, ethanol, ether, ethyl acetate, methanol, and chloroform) were employed to obtain extracts from leaves or seeds, resulting in markedly different antileishmanial activities and selectivity indices (Table 1). For instance, among the crude extracts from leaves, SI values ranged from as low as 2.32 (ethyl acetate extract) to as high as 64.8 (methanol extract), depending on the solvent used. The essential oil fraction obtained with ethanol exhibited a SI value of 110.0, further underscoring the critical influence of extraction method and fractionation on bioactivity outcomes. This variability can be attributed to the distinct chemical affinities of each solvent. Ethanol, particularly at intermediate concentrations (70–80%), allows the solubilization of a broad spectrum of secondary metabolites with medium to high polarity. Under optimized extraction conditions—such as reflux or ultrasound-assisted extraction—ethanol significantly enhances the recovery of phenolic compounds and flavonoids, which have been widely associated with antioxidant, anti-inflammatory, antimicrobial, and antitumor activities [20, 21, 22]. In contrast, highly polar glycosylated metabolites are more efficiently extracted with water, although this non-selective solvent also dissolves hydrophilic impurities that may affect bioactivity measurements. Solvents such as dichloromethane and ethanol tend to extract mid-polarity bioactive compounds like alkaloids, terpenes, and phenolics, while ether and ethyl acetate show similar but variable affinities depending on compound structure. Methanol extracts, which concentrate moderately polar and often lipophilic secondary metabolites, have been repeatedly associated with anti-Leishmania effects [23, 24, 25]. In contrast, non-polar solvents such as hexane are more likely to extract lipophilic substances like waxes and chlorophylls, which are generally considered non-bioactive in antiprotozoal assays [26] Taken together, these findings illustrate the critical influence of solvent selection on the phytochemical profile and biological performance of medicinal plant extracts and reinforce the importance of rational extraction strategies in antiprotozoal drug discovery.

Another noteworthy finding of this review is the high proportion of studies focused on *L. panamensis*, a species with broad geographic distribution and clear clinical relevance in Central America and along the Pacific coast of Colombia and Ecuador. Although this focus was expected given the local epidemiology, its confirmation is particularly significant in light of the global research landscape. Previous reviews have been conducted in regions such as East Africa (e.g., Ethiopia) and Asia (e.g., Iran), and have focused on *Leishmania* species that are not representative of Colombian transmission patterns: *L. tropica*, *L. major*, and *L. infantum* in Iran, and *L. aethiopica*, *L. donovani*, and *L. major* in Ethiopia. Besides *X. discreta*, we found that other Colombian plant species exhibited promising selectivity against *L. panamensis*, including *Origanum vulgare* (SI = 12.90) against promastigotes, and *Moquilea salicifolia* (SI >20.40), *Picramnia gracilis* (SI >11.80), and *Zanthoxylum schreberi* (SI = 11.60) against amastigotes. In the case of *L. braziliensis*, *Handroanthus chrysanthus* exhibited high selectivity (SI = 18.90, amastigotes), while for *L. major*, selective activity was observed in *Z. schreberi* (SI = 10.50, amastigotes) and *Ocotea macrophylla* (SI = 10.36, promastigotes). These findings support the notion that a subset of Colombian medicinal plants possess selective antileishmanial activity not only against the most prevalent local species but also against others of broader epidemiological relevance, thereby reinforcing their potential value as starting points for drug development beyond the local setting.

A major limitation of this review was the substantial methodological heterogeneity among the included studies. Some of this variability was expected, arising from differences in the plant parts used, extraction methods, presence or absence of fractionation, and the developmental stage of the parasite targeted (e.g., promastigotes, intracellular amastigotes, or axenic amastigotes). However, additional heterogeneity stemmed from the lack of standardized protocols, particularly in the methods and criteria used to evaluate antileishmanial activity. This inconsistency hindered meaningful cross-study comparisons and contributed to the high level of statistical heterogeneity observed. Furthermore, most studies lacked *in vivo* validation, limiting the translational potential of *in vitro* findings to more advanced preclinical models. Similar challenges have been reported in reviews conducted in other geographical regions, suggesting that this is a broader, structural issue within the field. Therefore, there is a pressing need to establish harmonized experimental guidelines for antileishmanial bioassays. These should include the systematic use of standardized positive controls, generation of complete dose-response curves, and parallel evaluation of parameters such as cytotoxicity in host cells. Adoption of these practices would significantly improve the quality, reproducibility, and translational relevance of the evidence generated.

## Conclusion

This systematic review and meta-analysis provides evidence that several medicinal plants collected in Colombia exhibit promising *in vitro* antileishmanial activity, with IC_50_ and selectivity index (SI) values that support their potential for further pharmacological development. The predominance of studies targeting *L. panamensis* aligns well with Colombia’s epidemiological context. *Xylopia discreta* emerged as a particularly compelling candidate against *L. panamensis*, showing consistently high SI values across multiple extraction methods. Other Colombian plant species that exhibited promising selectivity against *L. panamensis* include *Origanum vulgare*, *Moquilea salicifolia*, *Picramnia gracilis*, and *Zanthoxylum schreberi*.

Importantly, one of the key contributions of this study is the inclusion and systematic analysis of SI, an essential parameter for estimating the relative safety of plant extracts in host cells. While IC_50_ values reflect biological potency, the SI provides critical insight into the therapeutic window of a compound. Extracts with moderate IC_50_ but high SI may represent safer and more promising candidates than highly potent but cytotoxic ones. Therefore, incorporating SI into future evaluations is crucial for the accurate selection of leads for preclinical development.

In addition, differences in solvent polarity and extraction techniques were associated with considerable variation in both IC_50_ and SI values. This highlights the significant impact of extraction solvents on bioactivity outcomes, reinforcing the importance of rational extraction strategies in natural product research.

Despite these promising findings, the high methodological heterogeneity among studies and the frequent absence of *in vivo* validation limit the comparability and translational applicability of the available data. Variability in plant parts used, extraction protocols, parasite developmental stages, and outcome measures all contributed to the high statistical heterogeneity observed. To address these challenges, the establishment of standardized experimental guidelines, including the use of full dose–response curves, standardized positive controls, and cytotoxicity assays in host cells, is essential. The adoption of such practices would enhance the quality, reproducibility, and translational relevance of future antileishmanial research.

## CRediT authorship contribution statement

**CNC:** Conceptualization, Methodology, Validation, Formal analysis, Investigation, Data curation, Writing – original draft, Writing – review & editing, Visualization. **LM:** Conceptualization, Methodology, Formal analysis, Investigation, Writing – original draft, Writing – review & editing, Visualization. **GZ, ADA, ZCR, EPC, DT:** Investigation, Writing – review & editing. **JCO:** Conceptualization, Methodology, Formal analysis, Investigation, Writing – original draft, Writing – review & editing, Visualization, Supervision, Project administration, Funding acquisition.

## Supporting information

Supplementary data to this article can be found online at

## Data availability

All relevant data are within the paper and its Supporting Information files.

